# Reproducible Correlations Of Corpus Callosum And Cingulum Generalized Fractional Anisotropy With Anxiety Ratings In Healthy Participants

**DOI:** 10.1101/2020.12.08.404608

**Authors:** Sergey Kartashov, Alina Tetereva, Olga Martynova

## Abstract

Diffusion tensor imaging revealed that trait anxiety predicts the microstructural properties of fiber tracts between the anterior cingulate cortex, prefrontal cortex, and amygdala. However, the whole-brain structural connectivity has been rarely reported in the non-clinical populations with marginal deviations in trait anxiety and substantial changes in state anxiety. This work is aimed to assess the correlation of state and trait anxiety inventory (STAI) scores and their one-day deviation with white matter connectivity at the whole-brain scale. 64-direction diffusion-weighted images were collected in 25 participants without prior complaints of excess anxiety and/or clinical history of anxiety disorders. The self-reported ranking of STAI was collected twice with 24-hour interval: one day before and several minutes before the MRI scanning. A correlation analysis between the generalized fractional anisotropy (GFA) and the STAI ratings was performed for all regions of the brain. According to the diffusion connectometry, the most reproducible positive correlation of GFA and anxiety scores was observed for corpus callosum. For both days of psychological assessment, the left cingulum GFA correlated negatively with State anxiety ratings, while the right cingulum GFA was strongly associated with Trait anxiety. Our results suggest that the density of the corpus callosum and bilateral cingulum tracts are associated with the individual level of anxiety.

## 1 Introduction

Many neurophysiological studies of anxiety are performed using magnetic resonance imaging (MRI). The main tool to study the functioning of the brain networks responsible for personal or situational anxiety is functional MRI (fMRI). For example, study of changes of variance and scale-free properties of blood oxygenation level-dependent (BOLD) signal between resting-state fMRI sessions and fear memory extinction may reflect the residual brain activity related to the recent experience (Tetereva et al., 2020).

Also, of great interest in studies of anxiety is its relationship with the structural features of the brain. Several strategies are used here. The first is the study of morphological features in high resolution structural images (Yin et al., 2020). A change in the volume of white or gray matter in certain regions of the brain can be associated with the features of their functioning. Accordingly, a relatively low volume may indirectly indicate a lower potential for the functioning of a particular area due to trauma, disease, or features of the ontogenetic development.

The second strategy is the study of structural connections of the brain, restored from diffusion MRI (dMRI). Fibers of white matter act as wires through which a nerve impulse is transmitted from one region to another. A decrease in the number of connections affects the ability to transmit information, resulting in a weak potential for the implementation of the relevant function (Li et al., 2014). However, the structural connection itself reflects only the presence or absence of brain connections, without displaying possible changes in additional derived parameters obtained from dMRI data (fractional anisotropy, radial diffusion, mean diffusion) also reflect white matter density, axon diameter and myelination. The totality of these data allows us to assess the difference of integrity of the brain structure between, for example, patients and healthy volunteers.

To date, many works have been published describing the relationship between indicators of personal and situational anxiety with the parameters of diffusion MRI (fractional anisotropy, radial diffusion) within specific areas of interest that do not cover the entire subject’s brain. As a rule, the following regions are distinguished: anterior thalamic radiation, caudate nucleus, cerebellar peduncles (superior, middle, inferior), dorsolateral prefrontal cortex, ventrolateral prefrontal cortex (VLPC), orbitofrontal prefrontal cortex, anterior cingulate cortex, posterior cingulate cortex, stria terminalis, cuneus, precuneus, amygdala, hippocampus, parahippocampus, and insula (Sengul et al., 2019). The possible contribution of other areas is not considered and seems unlikely.

To obtain more complete information, the authors decided to conduct a correlation analysis of the levels of trait and state anxiety and the generalized fractional anisotropy (GFA) parameter of the whole brain without excluding any regions. We proceeded from the assumption that different levels of trait and state anxiety in people may be associated with the variability of the features of the structural organization of the brain.

GFA derived from dMRI and believed to reflect fiber density, axonal diameter, and myelination in the white matter of the brain. Unlike fractional anisotropy (FA), GFA allows researchers to obtain more detailed information on multiangle diffusion (Tuch D. S. et al., 2004). For the accurate calculation of the GFA coefficient, a dMRI protocol with 64 directions of diffusion coding gradients was used. This is an advantage over most similar studies. This should help to improve the accuracy and reliability of the information received.

We chose the State-Trait Anxiety Inventory (STAI; Spielberger et al., 1983) to assess anxiety. The STAI is the most frequently used measure of trait and state anxiety in both psychological studies and clinical settings. The validity of STAI was proved by test-retest reliability coefficients, which have ranged from .65 to .75 over a 2-month interval (Spielberger et al., 1983). In our study, we asked volunteers without self-reported anxiety complaints to fill in STAI twice with a 1-day interval to check the reproducibility of GFA correlations with anxiety ratings, which might insignificantly change within 24 hours.

## 2 Methods and materials

### 2.1 Participants and procedure

A total of 28 healthy right-handed volunteers (24+-4.4 years old, 10 females) without any neurological or psychiatric pathologies took part in the study. Before starting the MRI acquisition, each participant filled out a written informed consent form, an MRI safety questionnaire, and consent on personal data processing. The study protocol was approved by the local ethics committee of the NRC Kurchatov Institute, according to the requirements of the Helsinki Declaration. Data of three participants were excluded from the final sample due to MRI artifacts.

All participants completed the State-Trait Anxiety Inventory (STAI; Spielberger et al., 1970) twice: in one day before (Day 1: Trait1, State1) and on the same day (Day 2: Trait1, State1) of MRI-scanning session. Subjective ratings of State and Trait anxiety were separately subjected to analysis of variance for repeated measures (rsANOVA: within-subject factor - Day 1 vs Day 2) followed by Tukey post-hoc comparison.

The study was conducted in two days to assess the reproducibility of the anxiety level of each subject. If the levels of anxiety on the first and second days changed dramatically, then such people would be removed from the analysis.

### 2.2 Data acquisition

The scanning procedure was performed on an MRI scanner by Siemens Magnetom Verio 3T (Siemens Healthcare GmbH, Germany) in the Kurchatov’s Complex of NBICS nature-like technologies at the base of NRC Kurchatov Institute. The T1-weighted sagittal three-dimensional magnetization-prepared rapid gradient echo sequence was acquired with the following imaging parameters: 176 slices, TR = 1900 ms, TE = 2.19 ms, slice thickness = 1 mm, flip angle = 9°, inversion time = 900 ms, and FOV = 250 mm × 218 mm. A total of 25 diffusion MRI scans were included in the connectometry database. A DTI diffusion scheme was used, and a total of 64 diffusion sampling directions (TR = 13700 ms, TE = 101 ms, FoV = 240 mm × 240 mm, number of slices = 64) were acquired. The b-value was 1500 s/mm2. The in-plane resolution was 2 mm. The slice thickness was 2 mm. The b-table was checked by an automatic quality control routine to ensure its accuracy (Schilling et al., 2019). The diffusion data were reconstructed in the MNI space using q-space diffeomorphic reconstruction (Yeh et al., 2011) to obtain the spin distribution function (SDF). A diffusion sampling length ratio of 1.25 was used. The output resolution is 2 mm isotropic. The restricted diffusion was quantified using restricted diffusion imaging (Yeh et al., 2017). The generalized fractional anisotropy (GFA) values were used in the connectometry analysis.

### 2.3 Connectometry analysis

Connectometry analysis based on the “correlation tracking” paradigm and uses a nonparametric permutation test to determine the relationship of white matter pathways to anxiety level [5]. The result of the analysis is the pathway segments showing positive or negative correlation with the selected variable.

Diffusion MRI connectometry (Yeh et al., 2016) was used to derive the correlational tractography that has GFA correlated positively or negatively with Trait and State anxiety. A nonparametric Spearman correlation was used to derive the correlation. A total of 25 subjects were included in the analysis. A T-score threshold of 2.5 was assigned and tracked using a deterministic fiber tracking algorithm (Yeh et al., 2013) to obtain correlational tractography. The QA values were normalized. The tracks were filtered by topology-informed pruning (Yeh et al., 2019) with 2 iteration(s). An FDR threshold of 0.05 was used to select tracks. To estimate the false discovery rate, a total of 2000 randomized permutations were applied to the group label to obtain the null distribution of the track length.

## 3 Results

### 3.1 Anxiety assessment scores

The STAI scores were relatively in the moderate range: the average State Anxiety rating was 34.8+-11.43, the average Trait anxiety rating was 41.6+-12.31 at the initial day of filling the questionnaire. The range of anxiety fluctuated considerably: for State 20-71 (Fig S1, and for Trait 23-61 (Fig S2). However, the participants did not have any clinical manifestations of anxiety and were not under medical observation. As it was expected, State and Trait anxiety scores strongly correlated with each other (Spearman’ r = 0.69, p= 1.34e-4). There were no differences between both State and Trait anxiety ratings obtained at the first and second day of completing the questionnaires: F(1,27) = 0.62, p = 0.439 and F(1,27) = 3.44, p = 0.075, correspondingly. Test-retest reliability coefficients were 0.94 for Trait and 0.86 for State anxiety. We did not observe any significant correlation of STAI scores with gender or age of participants.

### 3.2 Correlations

The analysis revealed tractographic white matter pathways in which the GFA value had a positive or negative correlation with State or Trait anxiety score.

Fig1 shows pathways that correlated with Trait 1 positively (Corpus Callosum Forceps Major, Superior Longitudinal Fasciculus1 R, Corpus Callosum Tapetum, Corpus Callosum Body, Inferior Longitudinal Fasciculus R, Inferior Longitudinal Fasciculus L, Inferior Fronto Occipital Fasciculus R, Inferior Fronto Occipital Fasciculus L) and negatively (Cingulum Rarolfactory R, Superior Longitudinal Fasciculus1 R, Corpus Callosum Body, Inferior Fronto Occipital Fasciculus R, Corpus Callosum Forceps Minor, Cingulum Frontal Parietal R). Fig.2 shows pathways that correlated with Trait 2 positively (Corpus Callosum Forceps Major, Corpus Callosum Tapetum, Superior Longitudinal Fasciculus1 R, Anterior Commissure, Corpus Callosum Body, Inferior Longitudinal Fasciculus R, Inferior Fronto Occipital Fasciculus R, Inferior Longitudinal Fasciculus L) and negatively (Cingulum Rarolfactory R, Cingulum Frontal Parietal R, Fornix L, Cingulum Frontal Parietal L, Corpus Callosum Body). Fig.3 shows pathways that correlated with State 1 positively (Corpus Callosum Forceps Major, Corpus Callosum Tapetum, Superior Longitudinal Fasciculus1 R, Superior Longitudinal Fasciculus1 L, Middle Cerebellar Peduncle, Inferior Longitudinal Fasciculus R, Inferior Longitudinal Fasciculus L) and negatively (Cingulum Rarolfactory R, Cingulum Frontal Parietal L, Cingulum Frontal Parietal R, Fornix L). Fig.4 shows pathways that correlated with State 2 positively (Corpus Callosum Forceps Major, Corpus Callosum Tapetum, Corpus Callosum Body, Inferior Longitudinal Fasciculus R, Superior Longitudinal Fasciculus1 R) and negatively (Cingulum Rarolfactory R, Cingulum Frontal Parietal R, Cingulum Frontal Parietal L).

**Figure 1.**
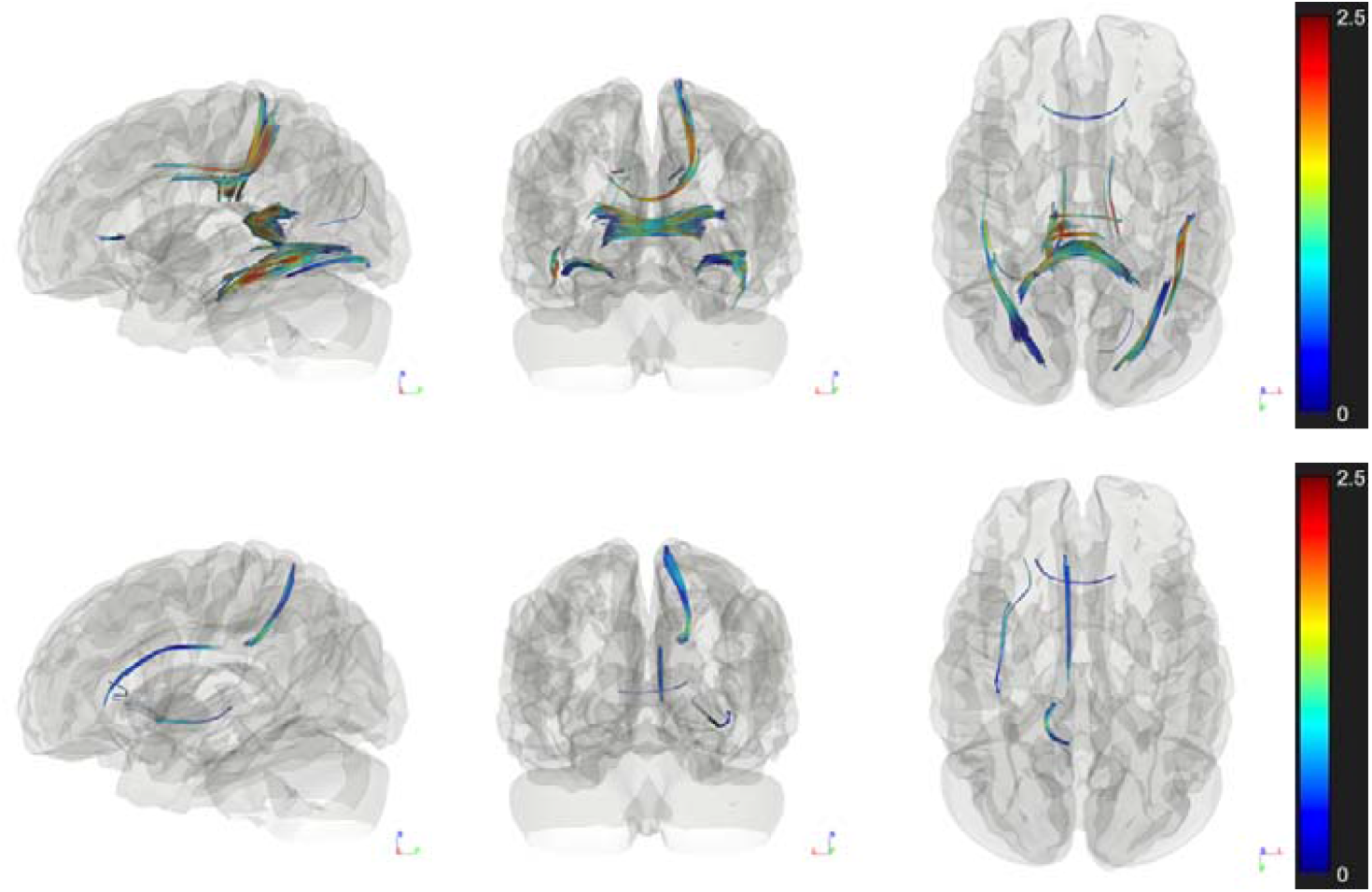
Diffusion MRI connectometry analysis shows a significant positive (top row) and negative (bottom row) correlation for Trait 1. T-score value is color coded.

**Figure 2.**
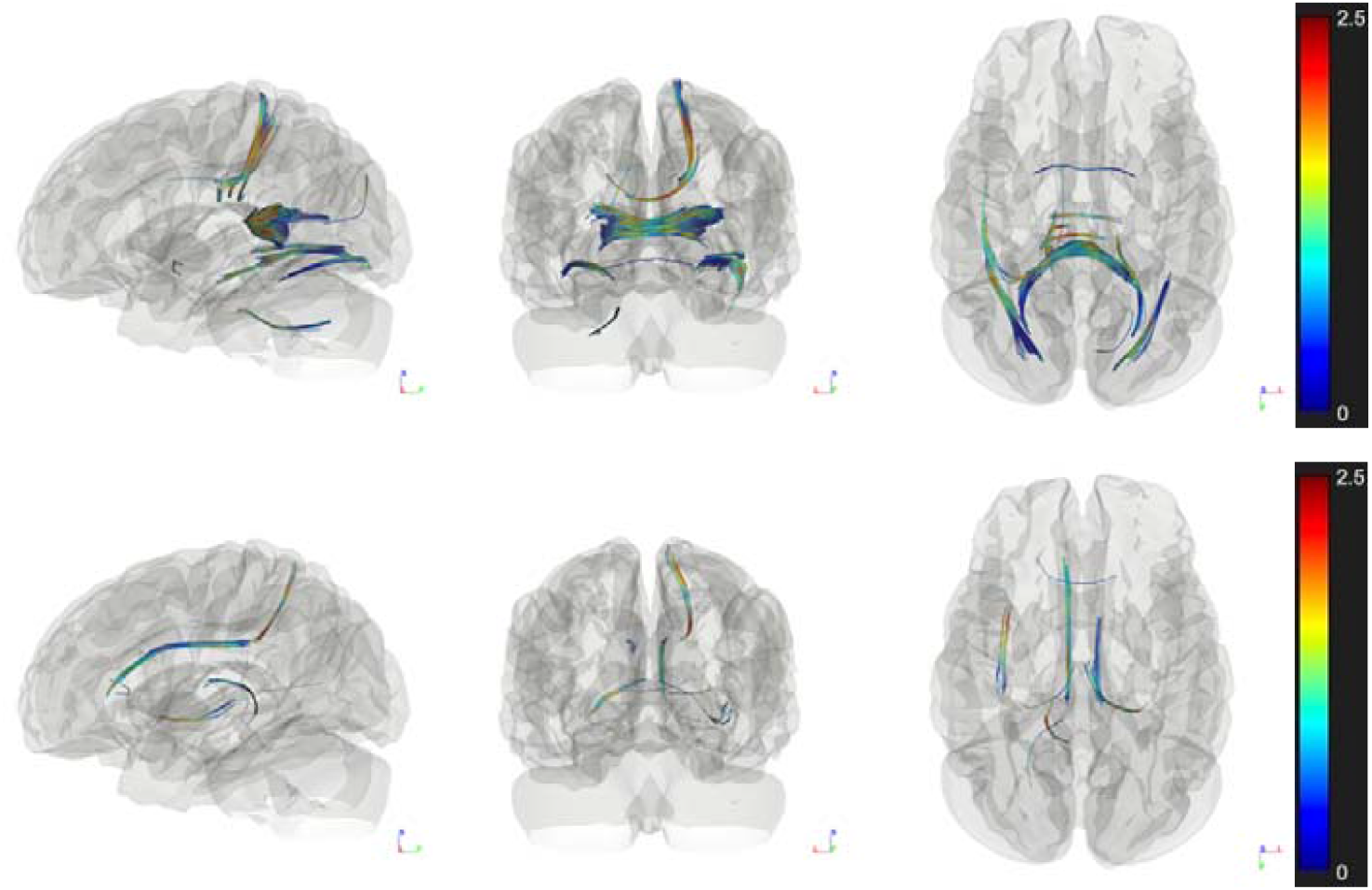
Diffusion MRI connectometry analysis shows a significant positive (top row) and negative (bottom row) correlation for Trait 2. T-score value is color coded.

**Figure 3.**
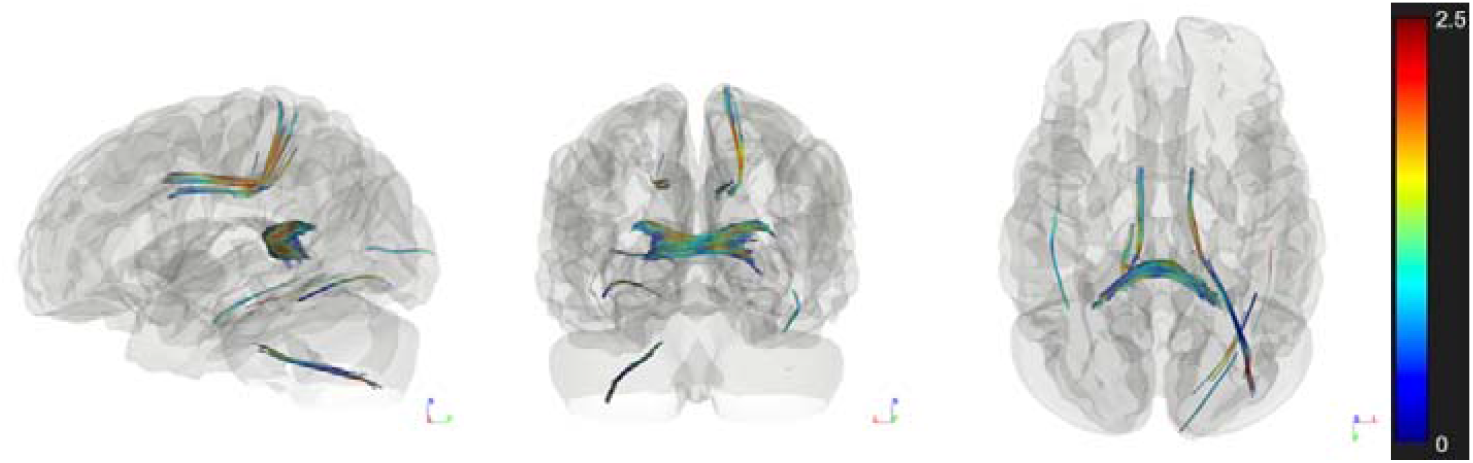
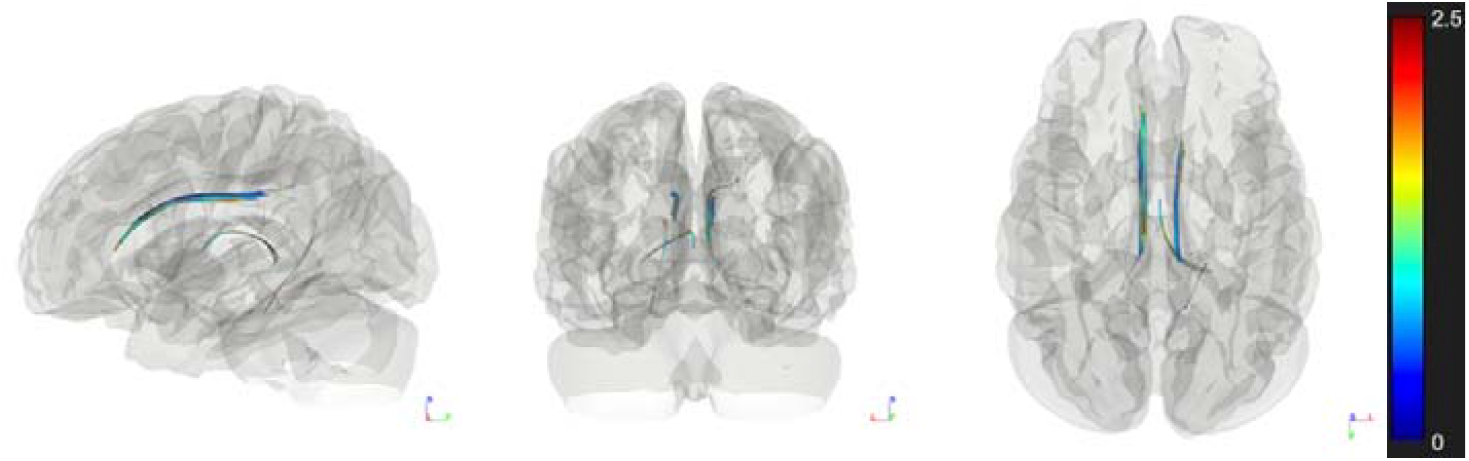
Diffusion MRI connectometry analysis shows a significant positive (top row) and negative (bottom row) correlation for State 1. T-score value is color coded.

**Figure 4.**
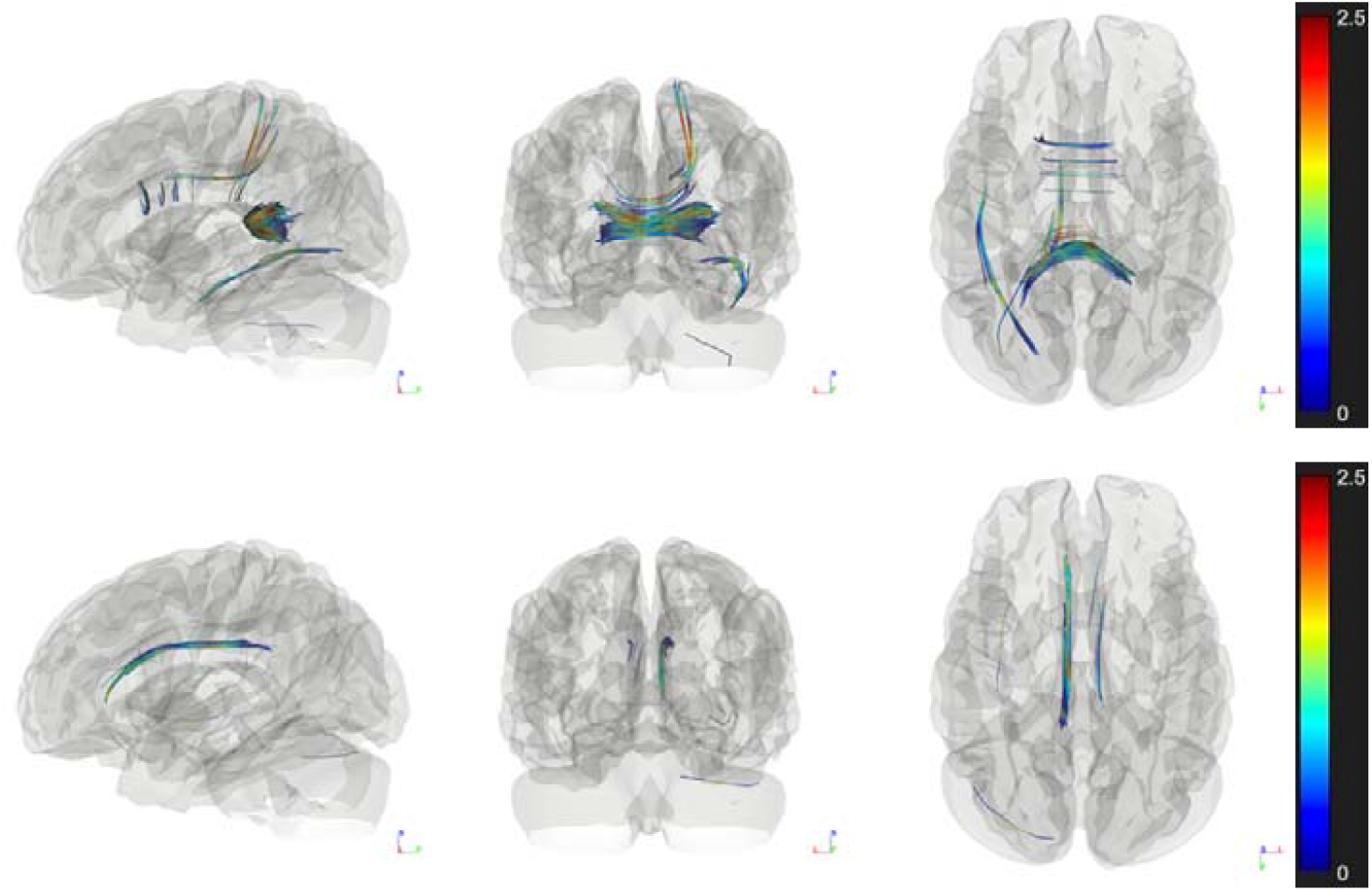
Diffusion MRI connectometry analysis shows a significant positive (top row) and negative (bottom row) correlation for State 2. T-score value is color coded.

**Table 1.**
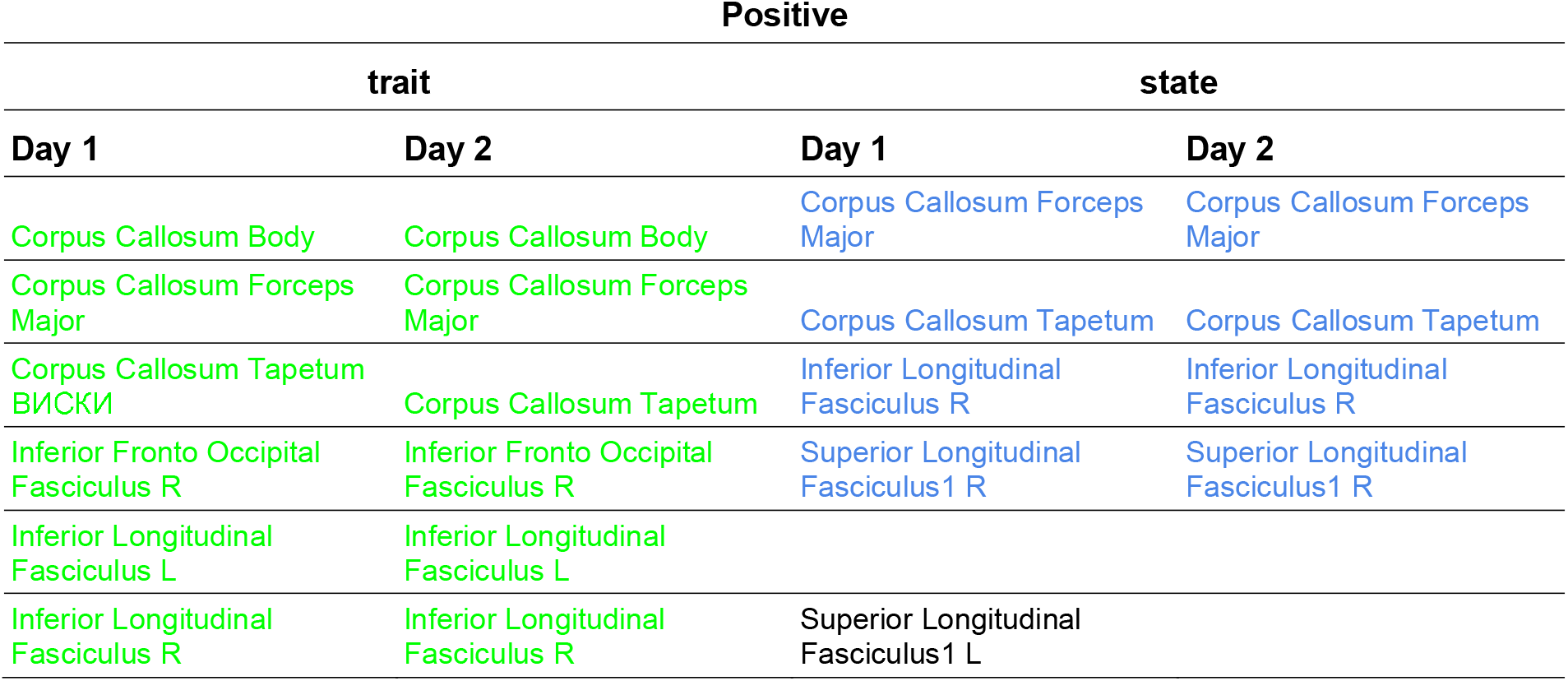
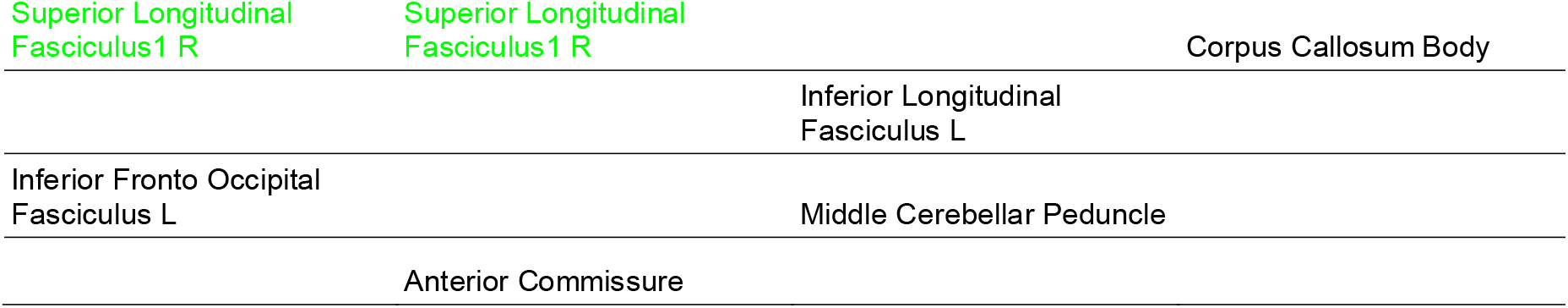
The positive correlations of generalized fractional anisotropy with Trait and State anxiety scores, obtained on Day 1 and Day 2 (the day of diffusion MRI acquisition). The same regions for the first and the second day are highlighted with color.

**Table 2.**
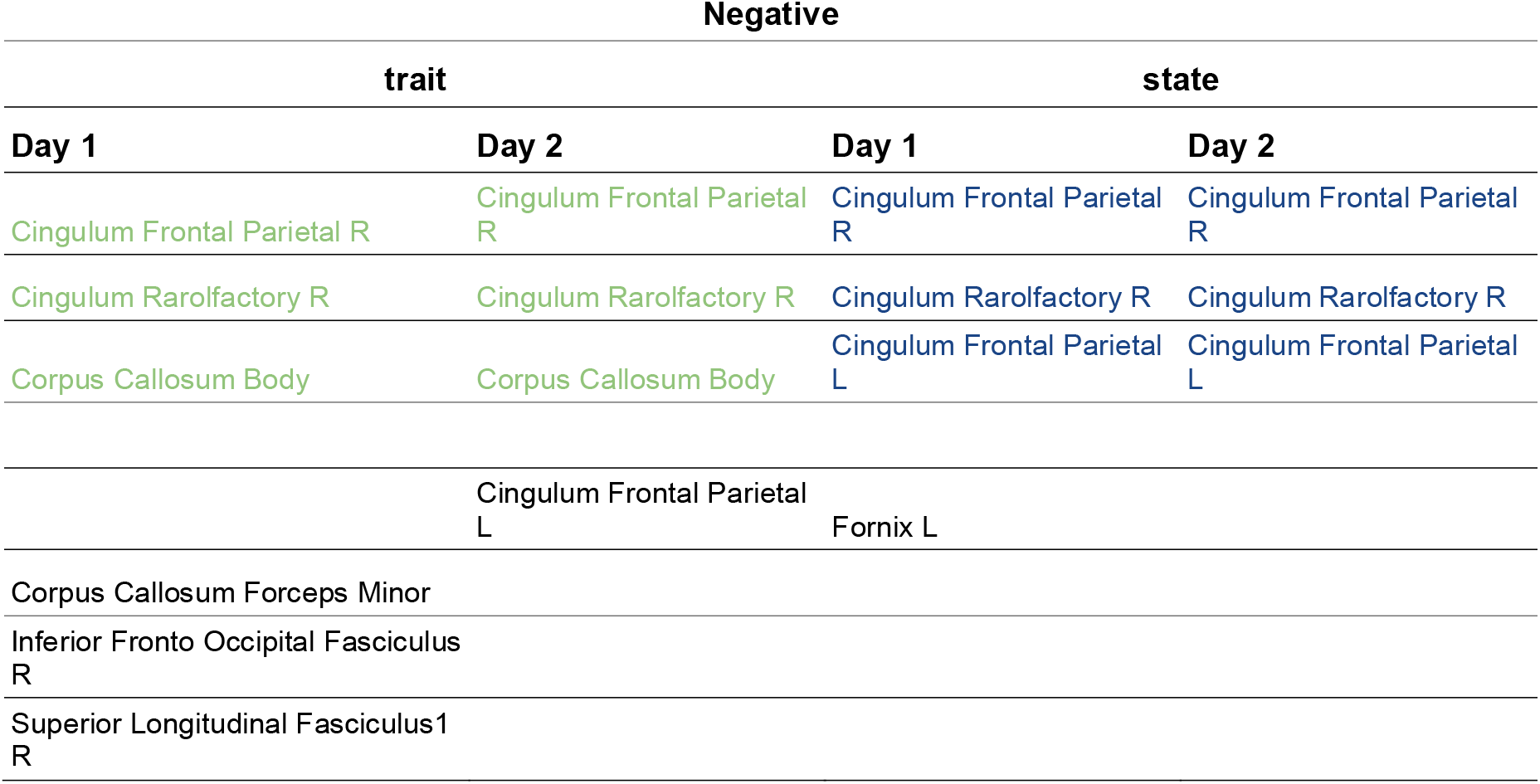
The negative correlations of generalized fractional anisotropy with Trait and State anxiety scores, obtained on Day 1 and Day 2 (the day of diffusion MRI acquisition). The same regions for the first and the second day are highlighted with color.

The analysis carried out for the first and second days indicates the reproducibility of the results, since the list of identified areas remained almost unchanged.

## 4 Discussion

Assessing the relationship between the structural organization of the brain and cognitive abilities in humans is one of the most interesting problems in neurosciences. Many research laboratories try to correlate behavioral responses with gray matter thickness in certain regions of the brain (Tuladhar et al., 2015), or with fractional anisotropy in the corpus callosum and other areas (Chiou et al., 2014), or with the number of connections between a limited number of regions of interest (Mišić et al., 2016). Thus, these works often do not analyze the entire brain as a whole.

In this article, the team of authors tried to slightly expand the search boundaries and performed Spearman’s correlation analysis for the Generalized Fractional Anisotropy (GFA) index among all voxels of the whole brain and information about the level of personal and situational anxiety.

The result of this analysis shows that there is a clear negative correlation of Trait and State Anxiety score with GFA in Right and Left Cingulum, respectively. Interestingly, Lochner et al. (2012) also found that lower diffusivity in the right dorsal cingulum was associated with increased anxiety and depression in OCD.

Either, according to the obtained data, Trait and State Anxiety levels are positively correlated with GFA in Corpus Callosum. However, in the work of colleagues (link https://www.ncbi.nlm.nih.gov/pmc/articles/PMC6406626/) such results are typical for depressive disorders.

In general, the search for neurophysiological correlates at certain levels of anxiety is further necessary to determine the pre-crisis state for serious depressive disorders.

