## Supplementary figures and images for "Reproducible Correlations Of Corpus Callosum And Cingulum Generalized Fractional Anisotropy With Anxiety Ratings In Healthy Participants"

### Figure S1

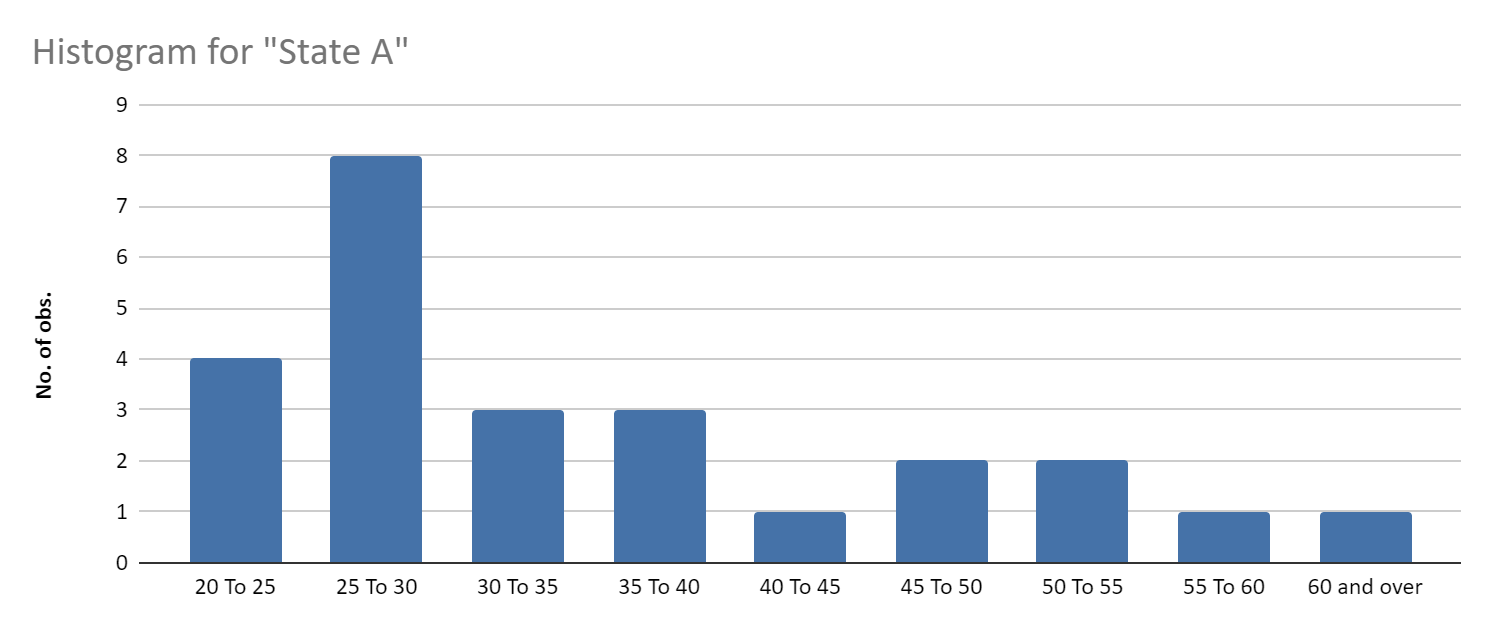

Figure S1. Histogram of State anxiety rating distribution among 28 participants.

### Figure S2

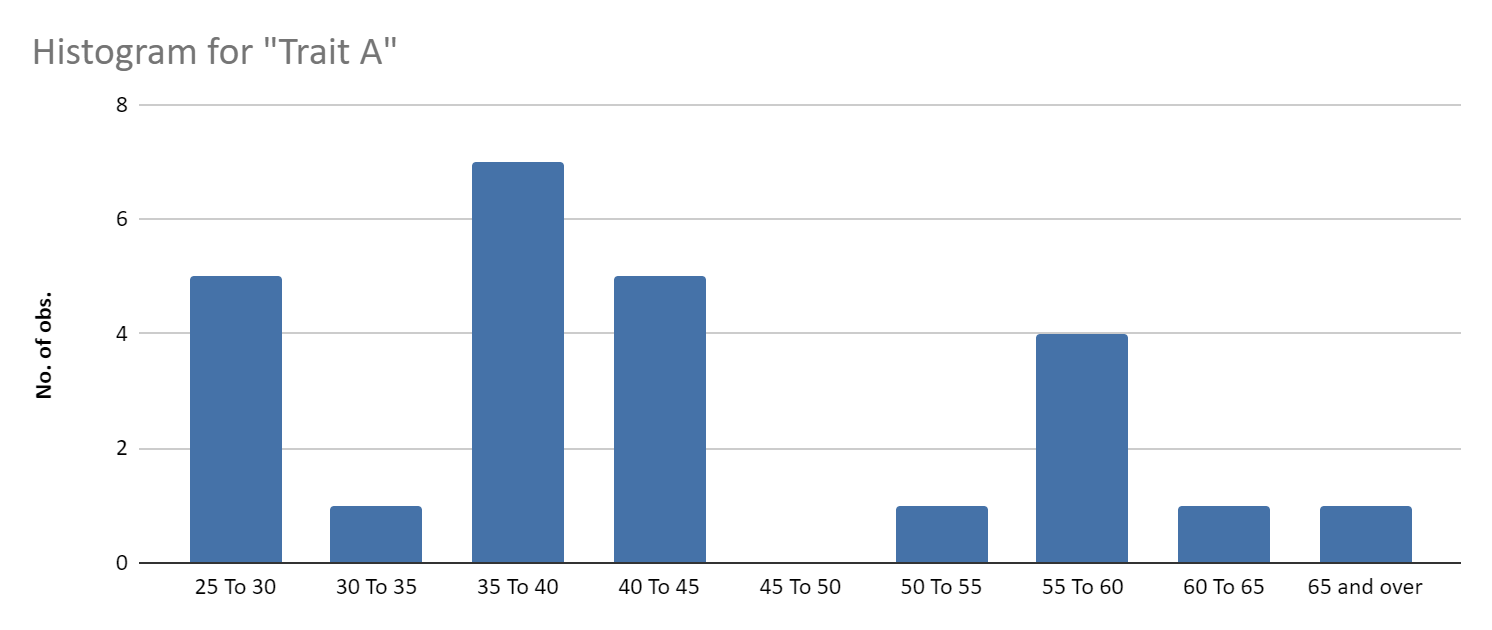

Figure S2. Histogram of Trait anxiety rating distribution among 28 participants.
